# Computational Model Predicts Paracrine and Intracellular Drivers of Fibroblast Phenotype After Myocardial Infarction

**DOI:** 10.1101/840017

**Authors:** Angela C. Zeigler, Anders R. Nelson, Anirudha S. Chandrabhatla, Olga Brazhkina, Jeffrey W. Holmes, Jeffrey J. Saucerman

## Abstract

The fibroblast is a key mediator of wound healing in the heart and other organs, yet how it integrates multiple time-dependent paracrine signals to control extracellular matrix synthesis has been difficult to study *in vivo*. Here, we extended a computational model to simulate the dynamics of fibroblast signaling and fibrosis after myocardial infarction in response to time-dependent data for nine paracrine stimuli. This computational model was validated against dynamic collagen expression and collagen area fraction data from post-infarction rat hearts. The model predicted that while many features of the fibroblast phenotype at inflammatory or maturation phases of healing could be recapitulated by single static paracrine stimuli (interleukin-1 and angiotensin-II, respectively), mimicking of the proliferative phase required paired stimuli (e.g. TGFβ and angiotensin-II). Virtual overexpression screens with static cytokine pairs and after myocardial infarction predicted phase-specific regulators of collagen expression. Several regulators increased (Smad3) or decreased (Smad7, protein kinase G) collagen expression specifically in the proliferative phase. NADPH oxidase overexpression sustained collagen expression from proliferative to maturation phases, driven by TGFβ and endothelin positive feedback loops. Interleukin-1 overexpression suppressed collagen via NFκB and BAMBI (BMP and activin membrane-bound inhibitor) incoherent feedforward loops, but it then later sustained collagen expression due to the TGFβ positive feedback loop. These model-based predictions reveal network mechanisms by which the dynamics of paracrine stimuli and interacting signaling pathways drive the progression of fibroblast phenotypes and fibrosis after myocardial infarction.

## Introduction

Wound healing is a complex process that involves a dynamic interplay between inflammatory and proliferative signaling. This process is especially important following injury to the heart, where cardiomyocytes are unable to regenerate. Scar formation and the preservation of viable heart muscle are important for continued cardiac function[1]. After myocardial infarction (MI), ACE inhibitors and beta blockers are prescribed to prevent adverse cardiac remodeling and heart failure [2], but the risk of heart failure and cardiac-related death post-MI remains high[3–5]. This is partly because wound healing is a balancing act between clearance of debris and formation of new scar, and the regulators of this dynamic process are not fully understood. Therefore, attempts to find therapeutic targets that allow for adequate collagen expression while avoiding excessive fibrosis have been largely unsuccessful. For example, although increased levels of interleukin-1 (IL1) have been linked to fibrosis and diminished cardiac index post-MI[6], blocking IL1 post-MI does not consistently improve healing and is actually associated with an increased risk of secondary MI[7, 8].

Myocardial infarcts follow a healing process that is similar to that in other organs [9, 10]. There is first an inflammatory phase characterized by extracellular matrix (ECM) breakdown and myocyte necrosis, which lasts around 2 days in rats and 5 days in larger mammals [11]. Then, the proliferative phase lasts around 2-5 days in rats (2 weeks in large mammals), during which fibroblasts proliferate, migrate into the wound, differentiate into myofibroblasts, and generate large amounts of collagen I and III and other ECM proteins [11, 12]. Ultimately, the wound matures into a stable scar with balanced ECM production and degradation. In the adult heart, cardiomyocytes do not proliferate sufficiently to re-populate the wound, so the ultimate fate of cardiac tissue depends on the behavior of cardiac fibroblasts. Excessive degradation can lead to ventricular dilation and wall rupture due to the loss of structural integrity in the heart wall [13]. Conversely, excessive ECM deposition, particularly in myocardium remote from the infarct, can lead to diastolic dysfunction [14, 15]. Many patients with heart failure post-MI have both dilation and fibrosis [1].

A beneficial infarct healing process likely involves a transient burst of collagen deposition that replaces lost cardiomyocytes with strong ECM without a sustained increase in ECM synthesis that leads to adverse remodeling [16]. This “transient fibrosis” is likely facilitated by many different factors including inflammatory cell phenotype and number, the pre-infarct signaling state, the size of the infarct, and the health of the remaining cardiac vessels [17, 18]. Fibroblasts play a prominent role throughout the entire wound healing process, and therefore present a good system for studying how cells respond to the dynamic signaling environment of wound healing[19]. Additionally, understanding how fibroblasts respond during the different phases of wound healing could identify mechanisms by which fibrosis develops in other organs.

Myocardial infarct healing is notoriously difficult to investigate because it involves many dynamic and interacting signaling processes. Fibroblasts are particularly difficult to study *in situ* during wound healing because they can differentiate from many different cell types and there is no clear consensus on fibroblast markers[20]. Computational modeling has been a useful method for investigating complex dynamic processes in many areas of biology. Although models have been constructed to study the wound healing process post-MI[21], no such model has yet been applied to study fibroblast intracellular signaling and phenotypic changes during myocardial wound healing[20, 22]. This study integrates a large-scale computational model of cardiac fibroblast signaling [23] together with post-MI experimental data for nine time-dependent paracrine stimuli to identify key paracrine and intracellular drivers of fibroblast phenotype after MI. This computational model of post-MI fibroblast dynamics was validated against independent experimental *in vivo* time courses of collagen mRNA and collagen area fraction throughout infarct healing. Next, we applied the computational model to identify simpler conditions with static single or paired paracrine stimuli that induce fibroblast phenotypes that mimic specific phases of post-MI healing. Virtual overexpression screens predicted both context-independent drivers of collagen synthesis as well as regulators that differentially affect collagen expression in the context of specific paracrine stimuli or phases of post-MI wound healing. Further model simulations generated experimentally testable mechanistic hypotheses for the signaling underlying such context-dependent responses, highlighting the utility of this computational model for interrogating the complex roles of fibroblasts in the post-MI wound healing environment.

## Methods

### Updates to Network Model of Fibroblast Signaling

We extended a large-scale computational model of fibroblast signaling [23] to be more relevant for subsequent experimental study of fibroblast phenotype dynamics *in vivo*. As described in previous studies[22, 50], the signaling model was implemented as a system of logic-based differential equations with default normalized reaction and node parameters. Here, the input nodes were separated from their associated ligand, and outputs associated with collagen maturation (e.g. LOX) and myofibroblast differentiation (e.g. contraction) were added[24–31]. See **Supplementary Methods** for more details. A schematic of the network model highlighting the added interactions and nodes is shown in **Figure S1**.

### Modeling the Dynamics of Post-MI Signaling and Collagen Expression

To simulate a post-MI setting, we fit idealized time-dependent curves to experimental timecourse data for the paracrine signals that act as inputs to the signaling model: IL1, IL6, tumor necrosis factor α (TNFα), angiotensin II (AngII), endothelin-1 (ET1), TGFβ, norepinephrine (NE), platelet-derived growth factor (PDGF), brain natriuretic peptide (BNP; corresponding node in the model is natriuretic peptide or NP). In the logic-based network model, inputs and signaling nodes are normalized to a continuous range from 0 to 1, where 1 represents a maximal level. The mechanical input was maintained at a normalized level of 0.6 throughout the post-MI simulation, because mechanical strain remains constant throughout the infarct healing timecourse in rats [32, 33]. Idealized time-dependent multi-exponential curves ranging from normalized levels of 0.1 to 0.6 were fit to timecoursedata from previous studies [34–41] (see **Supplementary Methods**). Due to a lack of available post-MI time course data from rat infarct zone alone, where needed, we incorporated studies that measured these paracrine factors in rat whole heart or border zone, or human serum post-MI (see **Table S1**).

To predict how single-cell changes in collagen expression lead to tissue-level changes in percent collagen area fraction, we adapted a previously published model of tissue-level collagen accumulation dynamics[42] and coupled it to the signaling model described above. The tissue-level model integrates time-dependent MMP activity and fibroblast numbers derived from previous studies of rat infarcts, as well as the collagen I and III mRNA levels predicted by the signaling model (**Figure S2**). MMP activity is approximated as a time-dependent function based on data from rat or mouse infarcts [43–46] rather than being defined by predictions from the signaling model, because MMPs are expressed by many cell types post-MI and fibroblast-specific expression does not fully account for cardiac MMP levels[47]. The timecourse of fibroblast numbers was also based on experimental data from rat infarcts[11]. This model is described in more detail in the **Supplementary Methods**.

We validated model predictions of post-MI collagen I and III mRNA dynamics against independent experimental data collected in rat infarcts [37, 48], in order to be consistent with the species used to develop the model and input curves (rat or human data). The predictions from the tissue-level model were validated against the collagen area fraction measured in rat infarcts [49].

### Modeling the Response to Static Paracrine Stimuli and Comparison with post-MI Dynamics

A constant stimulus of each single paracrine input and each input pair was set to a normalized level of 0.6, and the simulation was run to steady state. Steady-state predictions in response to static paracrine stimuli were compared to predictions at specific timepoints from the dynamic post-MI simulations (0 days, 1 day, 7 days, and 42 days) that correspond to different phases of infarct healing (pre-infarct, inflammatory, proliferative, and maturing respectively). We used principal component analysis (*pca* function in MATLAB) on mean-centered columns to compare static simulations and specific time points from the dynamic simulations based on output profiles that characterize the fibroblast phenotype (MMPs, collagens, other ECM proteins, proliferation, contraction, and αSMA). The single- and paired-paracrine static simulations that most closely matched the time points of interest from the dynamic post-MI simulation were identified by calculating their Euclidean distance within the space defined by the first three principal components.

### Screen for Intracellular Modulators of Collagen Expression

To identify modulators of collagen expression, we individually simulated overexpression of each node in the network and quantified its effect on the collagen expression nodes. These comprehensive screens were performed under conditions with static pairs of paracrine inputs representative of the different phases of post-MI healing, as well as with the time-dependent post-MI simulation that include nine dynamic paracrine stimuli. Overexpression of a given network node was simulated by increasing its normalized expression parameter (ymax) 10-fold. Change in collagen expression (the sum of activity from collagen I mRNA and collagen III mRNA) with overexpression (ymax = 10) was expressed as the difference from a simulation with no overexpression (ymax = 1) at the same timepoint and with the same paracrine stimuli.

## Results

### Expanding a fibroblast signaling network model to predict post-MI fibroblast phenotype dynamics

A published large-scale computational model of cardiac fibroblast signaling [23] was extended to make it more suited to predict the dynamics of fibroblast phenotype *in vivo* (see **Methods** and **Figure S1**). While the signaling model was originally developed and validated using a wealth of *in vitro* experimental data [23], we hypothesized that this model could be extended to predict post-MI fibroblast dynamics because it is capable of predicting semi-quantitative time-dependent behavior and it incorporates many of the pathways involved in infarct healing (e.g. IL1, IL6, TGFβ, AngII). Thus, we coupled the fibroblast network model with experimentally-based dynamic post-MI paracrine stimuli and a tissue-level model of collagen accumulation (**Figure 1A**).

**Figure 1:**
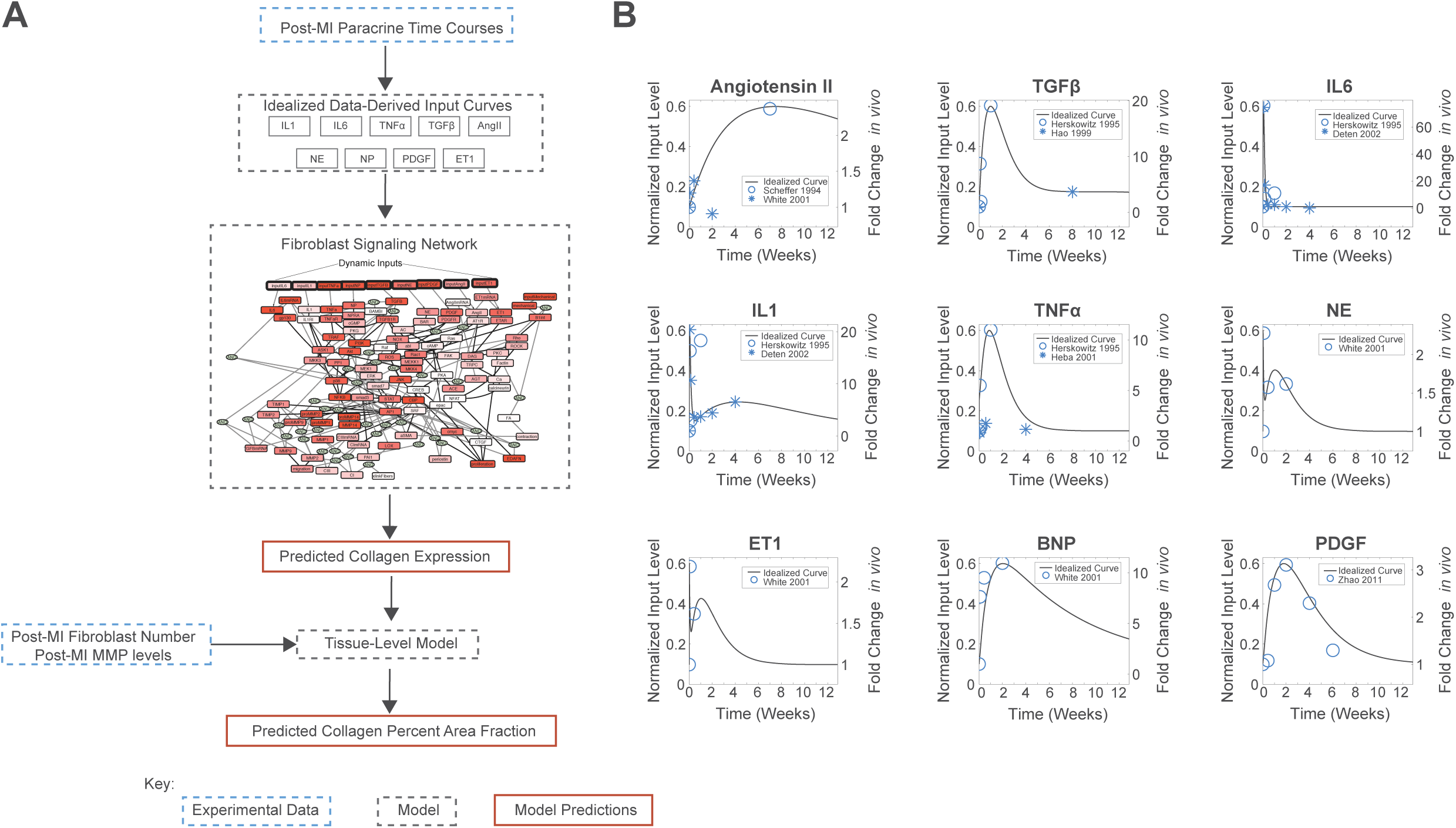
Computational model of post-MI fibroblast dynamics. A) Schematic of coupled model of post-MI fibroblast dynamics, incorporating dynamic paracrine stimuli, a fibroblast signaling network, and tissue-level collagen metabolism. B) Dynamic paracrine stimuli were modeled by fitting idealized multi-exponential curves to post-MI timecourse data from the literature [35–39, 41, 51]. These time-dependent signals provide inputs to the signaling network model. Experimental data were digitized from the indicated sources.

Timecourses for the nine paracrine model inputs were fit to multi-exponential curves based on published experimental data from rat or human following myocardial infarction (**Figure 1B**). Post-MI paracrine stimuli exhibited a range of distinct dynamic behaviors, including rapid transients (IL6), slower transients (e.g. TGFβ, PDGF), and rapid transients followed by a distinct slower phase (e.g. IL1, NE).

When driven by dynamic post-MI paracrine stimuli, the model predicts a wide range of distinct time-dependent network responses (**Figure S3**) due to the distinct behaviors of individual stimuli as well as pathway crosstalk and intra-network dynamics (**Figure S4** and **Video S1**). The model predicts transient collagen I and III mRNA expression dynamics that are similar to those observed in independent experimental data from rat infarcts (**Figure 2A**)[37, 48]. Further, the coupled network and tissue models predicted changes in collagen area fraction consistent with those measured in rat infarcts (**Figure 2B**)[32].

**Figure 2:**
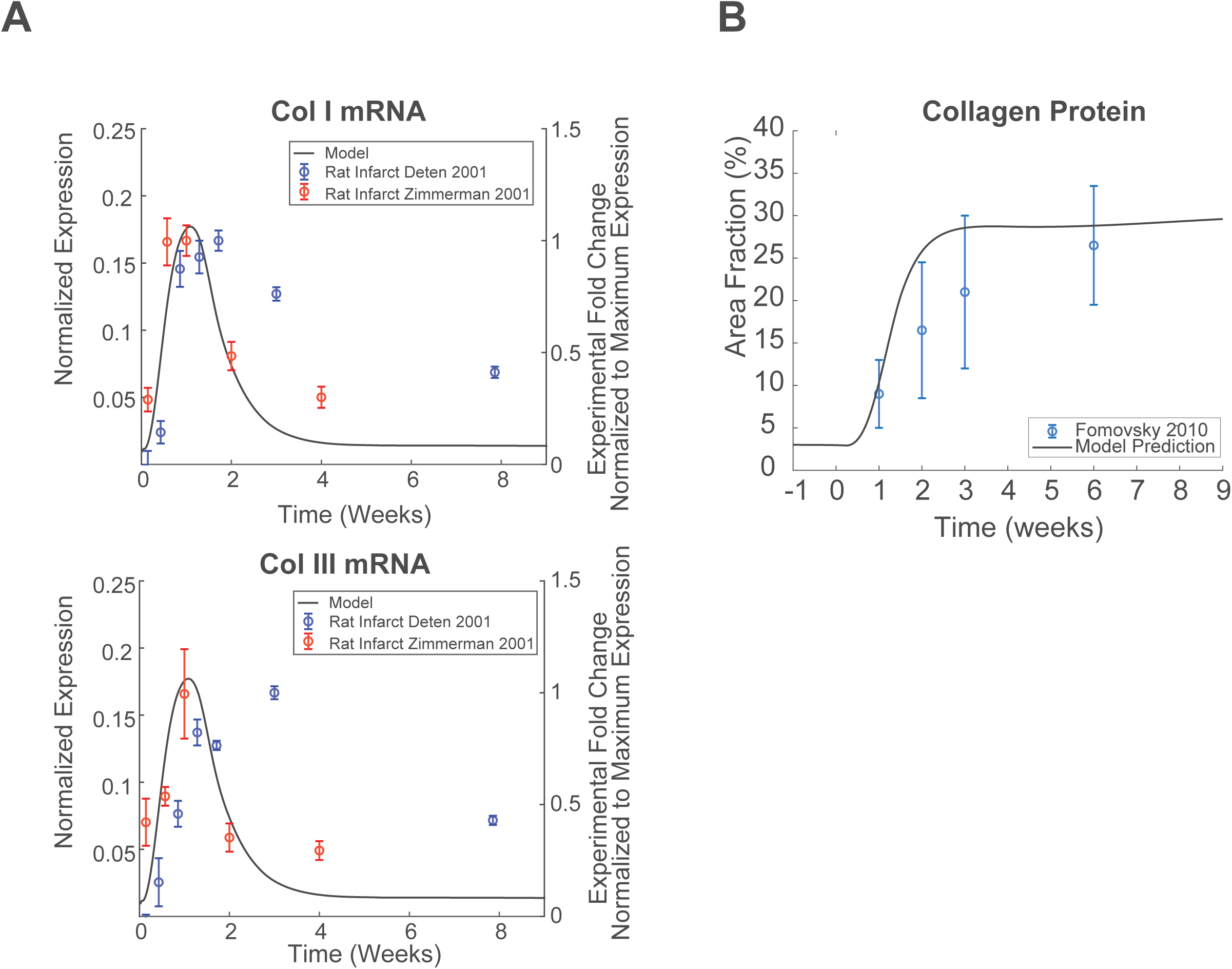
Modeling of post-MI fibroblast signaling reproduces dynamics of post-MI collagen expression and deposition. A) Validation of the predicted timing of collagen expression post MI against data from rat infarcts. B) Validation of predicted collagen accumulation (area fraction) post-MI from the tissue-level model. Experimental data were digitized from [37, 48].

### Relationship between fibroblast phenotypes induced by dynamic post-MI vs. static paracrine stimuli

*In vitro* experiments can provide precise control of simplified environmental conditions, but it is unclear to what extent such conditions can reproduce the phenotype of fibroblasts during the more dynamic and complex process of infarct healing[52]. Therefore, we used the model to identify individual or pairs of static paracrine stimuli that drive phenotypes that best mimic distinct phases of the post-MI fibroblast phenotype. The fibroblast model was run to steady-state under stimulation with static levels of the nine paracrine stimuli as well as all pairwise combinations (**Figure S5**). Using principal component analysis, we visualized the phenotypic relationship between fibroblasts stimulated by these 45 static paracrine conditions and the post-MI fibroblast phenotype timecourse (**Figure 3**). Of note, TGFβ stimulation had a very distinct effect on fibroblast phenotype, correlating with node loadings of pro-fibrotic outputs such as periostin, collagens, and αSMA near the negative PC2 axis of **Figure 3A** and **Figure 3B**. In contrast, the node loadings for MMP1/MMP14/proliferation and MMP2/MMP9 groups did not strongly associate with any single stimulus.

**Figure 3:**
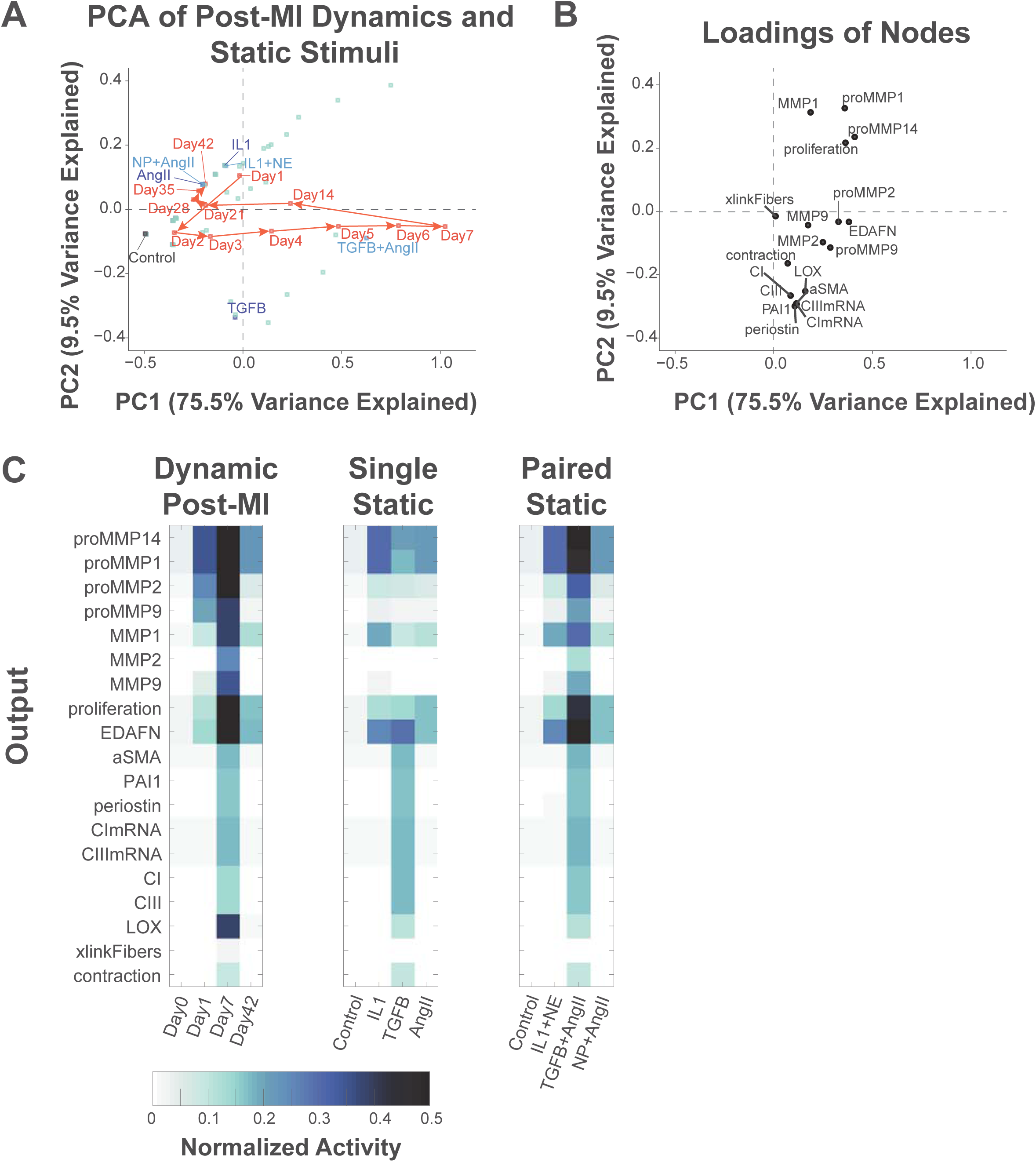
Simplified paracrine stimuli that mimic distinct phases of the post-MI fibroblast phenotype. A) Principal component analysis (PCA) to visualize the fibroblast phenotype at specific times post-MI (red circles) or at steady state with 45 static paracrine conditions (representative singles in dark blue, pairs in light blue). B) PCA node loadings show the contribution of each node towards the overall predicted fibroblast phenotype in the first two principal components. C) Activity profile of fibroblast phenotype nodes at selected timepoints from dynamic post-MI simulations (left), or at steady-state with the best matching single (center) or paired (right) paracrine stimuli.

We next examined the extent to which stimulation with one or two static paracrine factors mimicked fibroblast phenotypes at specific times post-MI. Such analysis can identify paracrine signals that drive specific phases of the post-MI fibroblast phenotype. The inflammatory phase (1 day post-MI) was fairly well mimicked by static IL1 (**Figure 3A**), which predicted proliferation, EDAFN, MMP1, and MMP14 expression but insufficient MMP2 or MMP9 expression (**Figure 3C**). The combination of IL1+NE improved predictions only slightly. In contrast, the period we defined as proliferative signaling at 7 days post-MI was not well mimicked by any single stimulus (**Figure 3A**), highlighting the signaling complexity of this phase (**Figure S3** and **Figure S4**). TGFβ mimicked aspects of the 7 day post-MI fibroblast phenotype (e.g. expression of collagens I/III, αSMA, periostin), consistent with previous studies showing that mouse fibroblasts at intermediate stages of post-MI healing are most similar to fibroblasts stimulated by TGFβ [52]. However, TGFβ was predicted to be insufficient to fully mimic the post-MI expression of multiple MMPs, proliferation, LOX, and collagen fiber cross-linking. Combination of TGFβ with AngII significantly improved predictions of MMP activity and fibroblast proliferation (**Figure 3C**). The late (42 day) time point was closely associated with AngII or the combination of NP+AngII, which mimicked sustained proliferation and expression of MMP1/MMP14. The paracrine pairs that induced phenotypes most similar to post-MI day 1, 7, and 42 fibroblasts are consistent with their elevation at those times post-MI as shown in **Figure 1**. Overall, these model predictions indicate that early and late stages of the post-MI fibroblast phenotype are more readily mimicked using single static stimuli, but intermediate post-MI timepoints involve many interacting stimuli and a complex network state that is more challenging to mimic *in vitro*.

### Predicting post-MI phase-specific regulators of fibroblast phenotype

Identification of regulators that drive distinct fibroblast phenotypes would provide a better understanding of the complex transitions that occur *in vivo* and may lead to potential phase-targeted therapeutics with reduced side effects. We simulated a virtual screen for regulators of collagen I/III mRNA expression by overexpressing each node in the context of paired paracrine static stimuli that best mimic the inflammatory (IL1+NE) or proliferative (TGFβ+AngII) post-MI phases, as well control conditions representative of the pre-infarct environment (**Figure 4**). For each node, overexpression was simulated by increasing that node’s normalized expression parameter 10-fold. Several nodes such as β1 integrin pathway members, ET1 pathway members, and MAP kinases like JNK were predicted to drive collagen expression across all three paracrine contexts.

**Figure 4:**
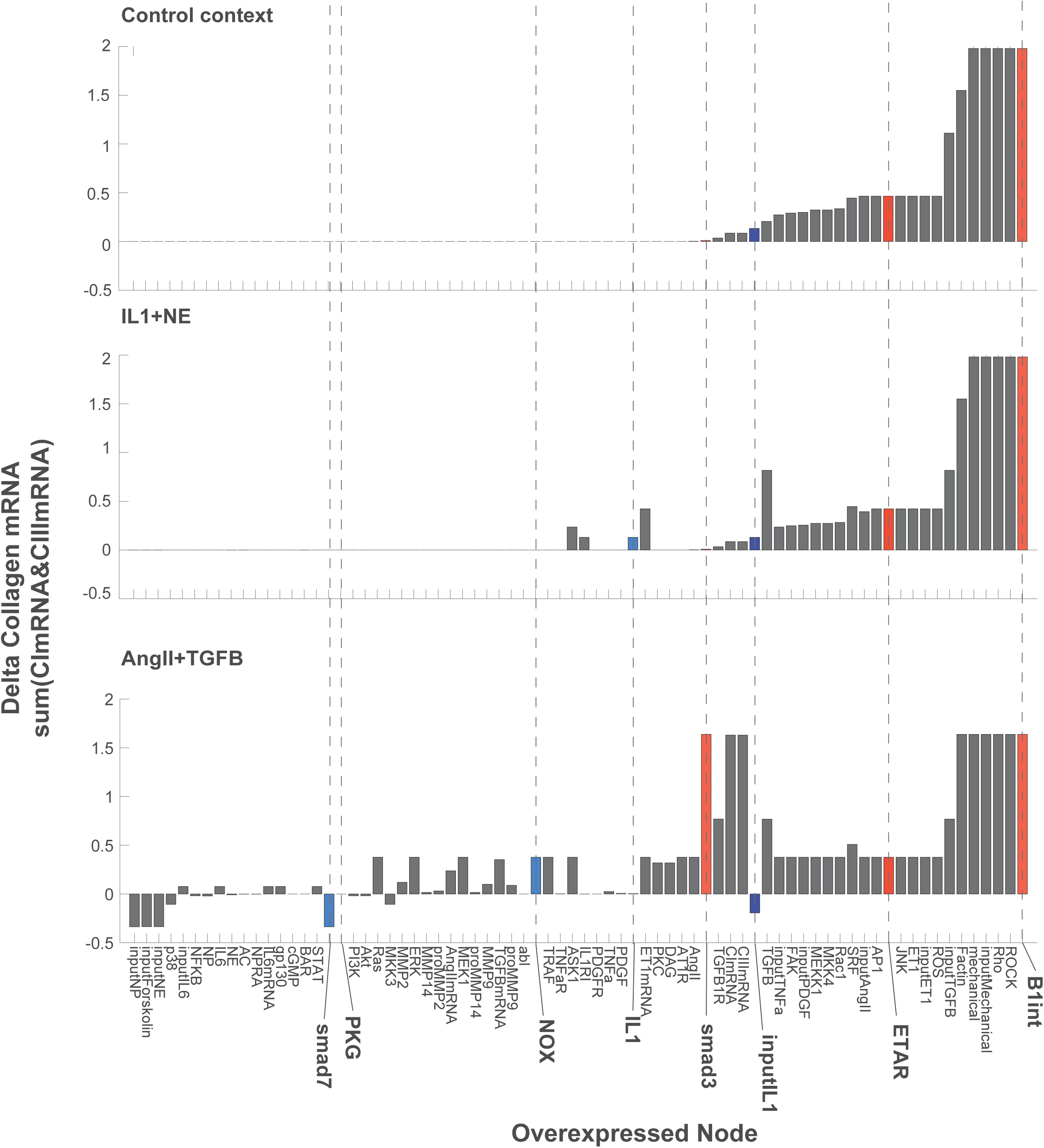
Modulators of collagen mRNA in the context of static paracrine stimuli that mimic inflammatory (IL1+NP) and proliferative (TGFβ+AngII) phases. Network nodes were each overexpressed 10-fold (normalized expression parameter ymax = 10) in the context of the indicated static paracrine stimuli (set to 0.6 normalized activity), predicting the change in collagen I and III mRNA compared to no overexpression. Nodes were rank-ordered by their predicted effect on collagen I and III mRNA expression with no paracrine stimulus (Control). Smad3, PKG, NOX, IL1, ETAR and B1int are emphasized for comparison with subsequent simulations of post-MI dynamics. Overexpressed nodes that did not affect collagen mRNA in any condition are not shown

The overexpression screen also identified a number of context-specific regulators of collagen expression. In the TGFβ+AngII stimulated condition, overexpression of TGFβ or AngII pathway members such as Smad3, ERK, or NOX enhanced collagen mRNA expression, while Smad7 and NP overexpression suppressed collagen mRNA. However, overexpression of these nodes did not significantly affect collagen mRNA expression in control or IL1+NE contexts. Overexpression of the “input IL1” node moderately increased collagen expression under control or IL1+NE stimulation but decreased collagen expression in the TGFβ+AngII context. This effect was dose-dependent, as overexpression of the “IL1” node itself did not sufficiently stimulate the IL1 receptor in the control and TGFβ+AngII conditions.

As static paracrine stimuli only partially mimic the history-dependent effect of the dynamic post-MI paracrine environment (**Figure 3**), it is unknown to what extent intracellular regulators of fibroblast phenotype identified under simplified conditions *in vitro* can explain their role *in vivo*. To identify post-MI phase-specific regulators, we performed a virtual overexpression screen of the 106 network nodes in the context of the nine dynamic post-MI paracrine stimuli. Activity timecourses for each of the overexpressed nodes in each of these 106 simulations are shown in **Figure S6**. Node overexpression induced a diverse range of phase-specific effects on the post-MI timecourses of collagen I/III mRNA expression (**Figure 5)**. Overexpression of some nodes was predicted to induce sustained collagen mRNA expression even before MI, such as TGFβ, ETAR, JNK, and β1-integrin pathway members. These context-independent nodes had an effect on collagen expression similar to that identified under static control conditions (**Figure 4**). In contrast, several regulators strongly increased (Smad3) or decreased (PKG) collagen mRNA only in the proliferative phase. Some regulators such as NOX not only increased collagen mRNA in the proliferative phase but sustained it into late phases. Other regulators had dichotomous effects, such as IL1, which enhanced collagen mRNA during the inflammatory and late phases but decreased collagen during the proliferative phase. Conversely, NFκB, IL6 and Akt overexpression modestly suppressed early and late collagen mRNA expression but enhanced collagen mRNA expression at intermediate timepoints.

**Figure 5:**
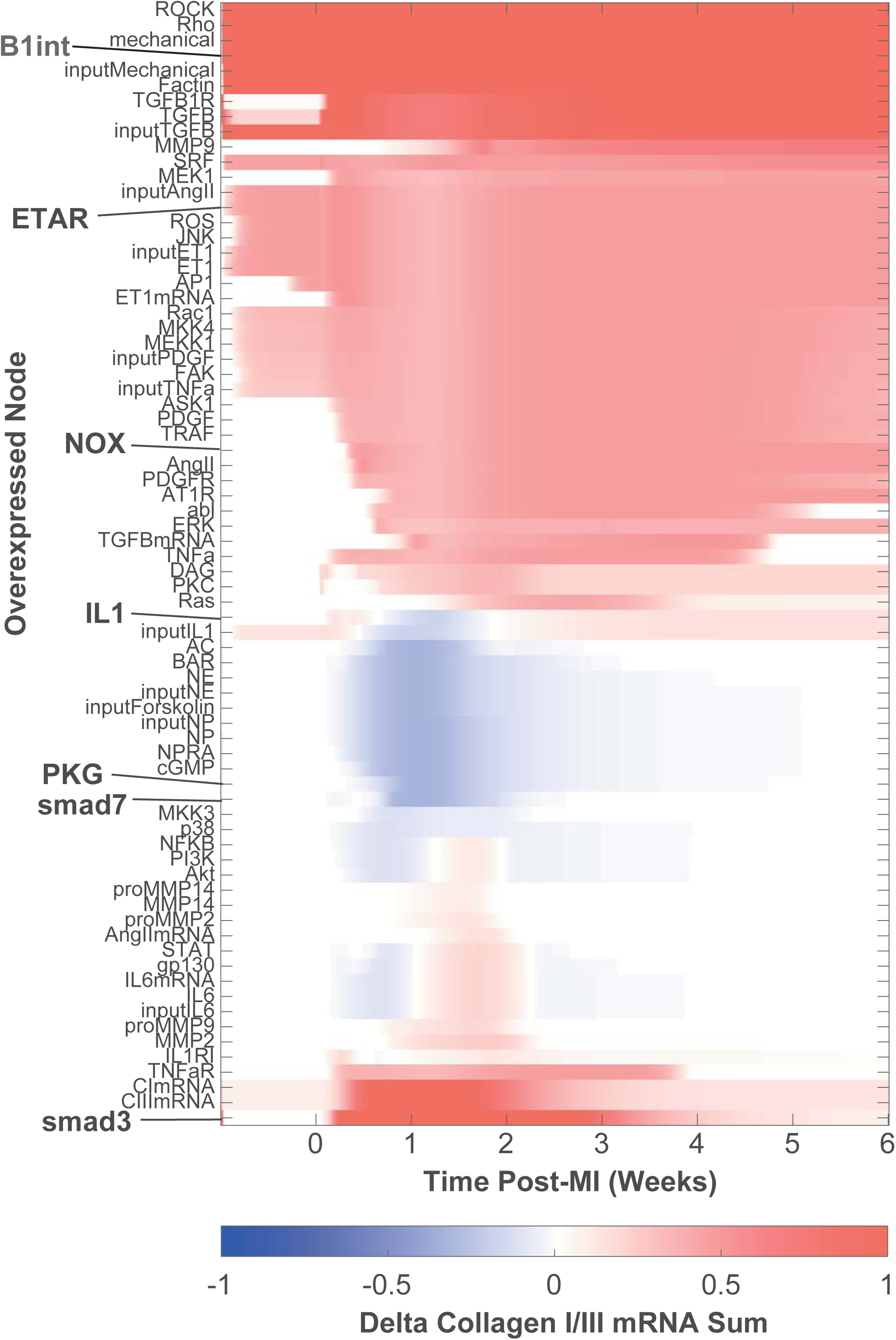
Post-MI overexpression screen to identify phase-specific regulators of collagen mRNA expression. Each row shows the effect of 10-fold overexpression of the indicated node. The predicted change in collagen expression is calculated as [Collagen I mRNA + Collagen III mRNA]_overexpressed_ - [Collagen I mRNA + Collagen III mRNA]_control_. Overexpressed nodes that did not affect collagen mRNA in any condition are not shown.

### Mechanisms contributing to post-MI phase-specific regulation of collagen

The mechanistic nature of the computational model enables identification of the specific conditions and network mechanisms that underlie phase-specific regulation of collagen expression. For specific influential nodes of interest, we performed additional simulations representing different levels of overexpression or knockdown using the dynamic post-MI model. We used the model’s mechanistic network structure along with sensitivity analysis as described previously [23] (data not shown) to identify simplified network mechanisms that mediate the effects of these notable regulators of collagen expression. For example, the model predicted that overexpression of Smad7 amplifies its activity and decreases collagen mRNA expression only in the proliferative phase (1-2 weeks), resulting in decreased collagen area fraction in the infarct throughout post-MI remodeling (**Figure 6A**). This prediction is consistent with in vitro studies showing Smad7 overexpression decreases collagen expression in vitro[53], as well as our simulations of the simplified AngII+TGFβ context (**Figure 3**).

**Figure 6:**
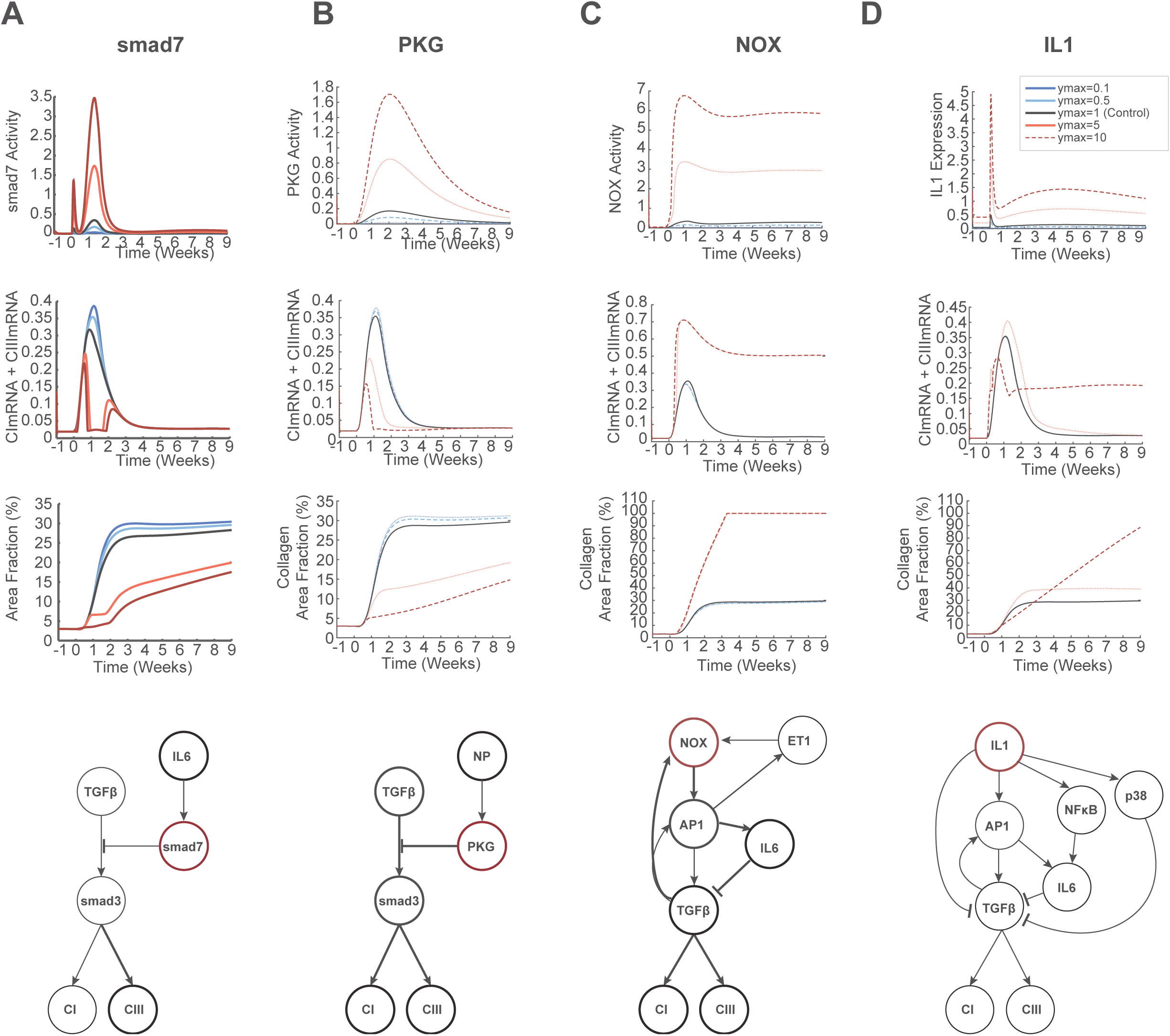
Mechanisms contributing to context-dependent regulators of collagen expression post-MI. Overexpression or knockdown were simulated by increasing or decreasing the normalized expression parameter (ymax) for Smad7 (panel A), PKG (B), NOX (C), IL1 (D). The resulting post-MI dynamics of that node’s activity, collagen mRNA expression, and collagen area fraction are shown. Simplified schematics indicate the network mechanisms by which these nodes regulate collagen expression.

Like Smad7, PKG was predicted to reduce collagen expression by inhibiting Smad3 activity during the proliferative phase. However, its inhibitory effect on collagen mRNA was more sustained due its more prolonged stimulation by the upstream paracrine NP (**Figure 6B**). Interestingly, PKG was not predicted as a negative regulator in the static AngII+TGFβ context context (**Figure 4**), as the simplified AngII+TGFβ context lacked the NP stimulation of PKG activity (**Figure S5**). In contrast to Smad7 and PKG, Smad3 overexpression strongly enhanced collagen mRNA in the proliferative phase with subsequent and sustained increase in collagen area fraction (**Figure S7**).

NOX overexpression was predicted to induce sustained NOX activity beginning in the proliferative phase post-MI, primarily due to activation of multiple positive feedbacks by AP1-TGFβ, NOX1-AP1-TGFβ, and AP1-ET1. The predicted AP1-TGFβ feedback involves both AP1-dependent transcription and MMP-induced activation of latent TGFβ. Together, these feedbacks induced sustained collagen I/III mRNA expression (**Figure 6C**). NOX also induced partial negative regulation of collagen mRNA via an incoherent feedforward circuit involving AP1-driven expression of IL6 and activation of Smad7. Overall, NOX overexpression induced a strong increase in collagen area fraction after MI.

IL1 exhibited a complex post-MI phase- and dose-dependent regulation of collagen expression. With moderate IL1 overexpression, collagen mRNA increased only moderately post-MI in the proliferation phase, driven primarily by the AP1-TGFβ positive feedback to produce a somewhat larger but stable infarct (**Figure 6D**). In contrast, strong IL1 overexpression enhanced collagen mRNA in the inflammatory phase, reduced collagen mRNA during the proliferative phase, and then subsequently provoked sustained collagen mRNA expression. The model predicted that the IL1 pathway has multiple mechanisms by which it negatively regulates collagen expression in the proliferative phase including: NFκB stimulation of the IL6/Smad7 pathway, p38-mediated inhibition of ERK, and inhibition of the TGFβ pathway through upregulation of the decoy receptor BAMBI (BMP and activin membrane-bound inhibitor). However in later phases, the sustained collagen mRNA expression was primarily driven by continued activation of the AP1-TFGβ positive feedback loop, resulting in delayed but continually elevating fibrosis post-MI (**Figure 6D**). IL1 knockdown was not predicted to significantly alter post-MI collagen expression. These predictions are in line with studies that have shown IL1 inhibition is most effective in limiting collagen expression post-MI when levels of inflammatory cytokines are high[8].

Both moderate and strong overexpression of β1-integrin induced substantial fibrosis even before MI, consistent with its effect across all simplified paracrine contexts (**Figure 4, Figure S6**) and the continual mechanical stress in the beating heart. Moderate ET1AR overexpression induced strong and sustained collagen expression beginning in the inflammatory phase post-MI, while strong ET1AR overexpression induced fibrosis even in baseline conditions (**Figure S7**).

## 3.5 Discussion

Cardiac fibroblasts are central mediators of wound healing and cardiac function after myocardial infarction [16, 49, 54], yet the complexity of the dynamic in vivo paracrine environment and the fibroblast intracellular signaling network hinders therapeutic targeting [55]. Here, we extended a large-scale computational model of the fibroblast signaling network to identify paracrine and intracellular drivers of extracellular matrix synthesis in specific phases of post-infarct healing. By integrating experimentally-based post-MI dynamics of nine paracrine stimuli, the model accurately predicted the dynamics of collagen expression as validated against experimental data from rat infarcts. The model was applied to screen for drivers of collagen expression in the context of static paracrine stimuli representative of specific post-MI phases, as well as the more complex dynamic post-MI paracrine environment. These virtual overexpression screens identified post-MI phase-specific regulators such as PKG and Smad7, which suppress collagen expression in the proliferative phase via inhibition of Smad3; NOX, which sustained fibrosis in proliferative and late phases via AP1-TGFβ feedback; and IL1, which induced alternating enhancement and suppression of collagen expression. This study highlights how fibroblast signaling pathways coordinate to dynamically control matrix synthesis during wound healing, and invites further investigation into therapeutics that target fibroblasts at specific post-MI phases.

### Loci of control in the fibroblast signaling network

The complex, concomitant dynamics of many signaling pathways during infarct healing has made it difficult to experimentally identify the signaling mediators that drive collagen production[52]. Computational modeling enabled a detailed mechanistic investigation into the determinants of collagen production by integrating many sources of experimental data into a single framework. For example, the model predicted that Smad7 overexpression decreases collagen expression during the proliferative phase is consistent with previous studies that found decreased collagen expression in fibroblasts after overexpressing Smad7 *in vitro* [56] or after AngII-induced cardiac remodeling in mice [57]. The prediction that PKG overexpression reduces collagen expression post-MI is consistent with a previous study showing that upstream NP decreases post-MI fibrosis [58]. This literature-based fibroblast network model predicts that both Smad7 and PKG affect collagen expression via mechanisms that inhibit Smad3 [53, 59]. It is well-understood that TGFβ induces collagen production, but inhibiting TGFβ signaling before or after an MI can decrease the necessary collagen deposition during wound healing and increase the risk of cardiac dilation and mortality in rats[56]. Further, Smad3 knockdown was predicted to decrease post-MI fibrosis, consistent with previous studies[60]. Overall, these simulations indicate that Smad3 is a critical target for negative regulation of collagen expression in cardiac fibroblasts, which could direct future studies into other inhibitors of Smad3 activation as a way to modulate excessive fibrosis.

Inflammation has been extensively tested as a potential driver of cardiac fibrosis. It has been shown that higher pre-infarct levels of inflammation, high post-infarct levels of inflammatory mediators, and longer duration of inflammation are associated with an increase in fibrosis and a decrease in cardiac function post-MI[6, 8, 55, 61]. However, the use of broad anti-inflammatory treatments such as NSAIDs or corticosteroids as well as therapies targeted to specific inflammatory mediators have been shown to worsen outcomes rather than improve cardiac health post-MI[7, 13, 17, 55, 62–65]. Furthermore, although IL1 receptor inhibitors (e.g. Anakinra) have been shown to decrease cardiac fibrosis, they increase the risk of major cardiac events and heart failure long term[7, 63]. Sano et al recently showed that IL1 receptor inhibition attenuated post-MI fibrosis in mice with a TET2 mutation that increased inflammation, but in control mice, IL1 inhibition had no effect on collagen area fraction[8]. This is consistent with our prediction that strongly increasing IL1 signaling increases collagen synthesis long-term, while decreasing IL1 did not affect collagen. The role of IL1 in simultaneously inducing and attenuating TGFβ signaling has not been fully elucidated in the heart. *In vitro* studies of the effect of IL1 on fibroblast phenotype have not been fully consistent, highlighting the fact that IL1 might have a very context-specific role. We identified three mechanisms by which IL1 can attenuate TGFβ-induced collagen expression: the IL6 pathway, p38 signaling, and BAMBI upregulation. Examination of both bulk RNA-seq from fibroblasts post-MI and single-cell RNA-seq from Wnt-expressing fibroblasts post-MI shows increased expression of BAMBI in the proliferative phase [19, 66]. Further, BAMBI knockout enhanced profibrotic responses in TGFβ-stimulated cardiac fibroblasts or after transverse aortic constriction *in vivo* [67].

The model also predicted that NOX overexpression would increase collagen, while NOX knockdown would not affect collagen expression. However, decreasing NOX activity was previously shown to attenuate collagen expression post-MI[68]. This discrepancy between the model predictions and *in vivo* data is likely due to the model not predicting sufficient NOX activity post-MI without overexpression. NOX is downstream of TGFβ, ET1, and AngII, which both our model predictions and previous experiments demonstrate to enhance cardiac fibrosis[69, 70]. In the model, the mechanism by which both NOX and IL1 overexpression increase collagen expression is via an AP1-TGFβ positive feedback pathway. This predicted feedback loop is consistent with our previous modeling of mechano-activated fibroblasts validated in collagen gels, previous studies with lung fibroblasts, and in vivo studies demonstrating the critical role of AP1 component c-Jun in many fibrotic diseases[23, 71, 72]. Thus the role of AP1-TGFβ positive feedback could be the focus of further studies into potential anti-fibrotic therapies.

### The dynamic model as a tool for investigating wound healing

There is a need for computational tools for investigation of wound healing in order to more fully understand the development of fibrosis and identify potential therapeutic strategies against it[20]. In the heart, early collagen production is important for generating a strong scar[73], but continued production and fibrosis contribute to diastolic dysfunction[1, 16, 74]. It has been shown that the timing of collagen production is important in determining the ultimate health of the heart[64].

The dynamic modeling approach outlined in this study allowed for a direct comparison between specific signaling pathways and the phases of fibroblast phenotype, and this could be a tool for further investigation into cardiac fibrosis. We identified representative stimuli that mimic fibroblast activity in the pre-infarct (control), inflammatory (IL1+NE), proliferative (TGFB+AngII), and mature (NP+AngII) phases of wound healing. These stimuli can be used to design *in vitro* experiments mimic *in vivo* fibroblast phenotypes characteristic of specific time points during infarct healing. Future studies could investigate how changing the timing of these signals affect collagen expression. The model could be used to test the effect of potential novel therapeutics on fibroblast phenotype. The modeling of differential expression of nodes in the network invites the possibility of integrating gene expression data to investigate potential genetic causes of adverse wound healing that lead to heart failure, patient-specific variability, or single-cell heterogeneity. Furthermore, the model includes explicit network mechanisms, which guide *in vitro* experiments to validate promising collagen regulators identified in this study.

### Limitations and future directions

The main limitation of this study is that the model predicts post-MI contributions of fibroblasts but not other cell types such as macrophage and cardiomyocytes can alter cardiac remodeling. The tissue level model only indirectly accounts for the activity of these cells in collagen degradation and expression. Of note, the tissue level model does not account for changes in MMP expression by fibroblasts, as the tissue-level MMP activity that drives collagen degradation is assumed to be largely due to inflammatory cells. Furthermore, processes such as re-vascularization (which can improve infarct healing) or further cardiomyocyte injury (which can re-start or prolong wound healing and induce pathologic remodeling) are not captured by this model. As in many experimental screens and our past modeling of in vivo hypertrophy, here we performed virtual overexpression screens. This approach has advantages of testing whether overexpression is sufficient to modulate fibroblast phenotype and is more readily validated experimentally with transgenic mice or viral vectors. Future studies may more systematically explore simulations of genetic knockouts or small molecule inhibitors. However, this model is a useful step toward investigating additional details of fibroblast signaling dynamics and intercellular cross-talk post-MI.

In this study, we demonstrate that by integrating multiple time-dependent paracrine stimuli, the fibroblast network model can predict many features and regulators of post-MI fibroblast signaling and matrix remodeling. These predictions should be validated *in vivo*, but initial validations may be most practical *in vitro* with the static paracrine stimuli that we identified to mimic post-MI phases. Further studies could test how variations in the timing of input stimuli affect fibroblast activity or incorporate this model into a multi-scale model that can predict the effect of cell-cell interactions between macrophage and cardiomyocytes. Future studies will also be necessary to determine how therapeutic modulation of collagen dynamics can be leveraged to optimally improve cardiac remodeling and contractility after myocardial infarction.

### CRediT author statement

Angela Zeigler: Conceptualization, Writing - Original Draft, Software, Formal Analysis; Anders Nelson: Software, Formal Analysis; Anirudha Chandrabhatla: Software, Formal Analysis; Olga Brazhkina: Formal Analysis; Jeffrey Holmes: Conceptualization, Writing-Reviewing and Editing; Jeffrey Saucerman: Conceptualization, Writing-Reviewing and Editing.

## Acknowledgements

We thank Dr. William Richardson for valuable discussion of this work. This work was supported by the National Institutes of Health [grant numbers HL127944, HL137755, HL116449, HL007284]; the National Science Foundation [grant numbers 1560282, 1252854] and the University of Virginia Center for Engineering in Medicine. The funding sources had no involvement in the conduct of the research or decision to publish.

