## Supplementary Materials for "Computational Model Predicts Paracrine and Intracellular Drivers of Fibroblast Phenotype After Myocardial Infarction"

### Supplementary Methods

*Alterations to the original model*

Additions were made to the original fibroblast signaling network model [1] to make predictions more comprehensive of fibroblast phenotype and its effect on the extracellular matrix (see **Figure S1**). Specifically, we introduced three new nodes into the network: LOX, x-linked fibers (cross-linked collagen fibers), and contraction. The new reactions were supported by at least two published studies that used rat or human fibroblasts - the same criteria used in the initial development of the model network [1], with sources provided in **Table S1** [2–9]. Additionally, we separated the nodes that represent the levels of the exogenous paracrine inputs to the model from the nodes that represent the paracrine signals available to bind to each receptor, as indicated by the green nodes at the top of the network on **Figure S1**. This allowed the exogenous paracrine signals to be defined in a time-dependent manner as described below, while still modeling the autocrine behavior of some signaling inputs such as AngII. The form of the logic-based differential equations used to model these interactions and the default node parameters ($y_{initial}$, $y_{max}$, tau), reaction parameters (weight, Hill coefficient, EC50) are the same as those used in the original model.

*Input Levels*

Given the importance of fibroblasts in the post-MI wound healing process, we sought to leverage a computational model of fibroblast signaling [1]to model the effect of changes in cytokine and chemokines on fibroblast phenotype. This allows for a uniquely mechanistic study of post-MI signaling. Additionally, the methods described here outline a protocol for utilizing similar computational models to investigate dynamic signaling processes that are difficult to study *in vivo*.

Dynamic levels for all paracrine inputs were defined by curves based on measurements of those inputs in rat or human hearts post-MI. Measurements of IL1, IL6, TNFα, AngII, and PDGF were used from rat infarct where available or from rat peri-infarct tissue, whole heart extracts, or cardiac tissue remote from the infarct as indicated in **Table S2**. To our knowledge, there is no published post-MI rat study where NE, ET1, or NP were measured at multiple timepoints, so post-MI human data was used for those inputs.

For each input signal, multi-exponential curves were fit to the post-MI input data. Experimental data of paracrine inputs were first converted to fold-change by normalizing to the first timepoint. Next, we fit either a bi-exponential (**Equation 1**) or a sum of bi-exponentials (**Equation 2**) to the data, depending on the number of peaks indicated by time course data. The fitted input curves were then normalized to start at a control normalized level of 0.1 and reach a peak of 0.6 at the time corresponding to the peak signal observed experimentally. Curve parameters for time constants were manually fit to the kinetics of the experimental data, and the magnitude parameter ($p_1$ or $p_4$) was adjusted to ensure that the peak of the input curves was within $10^{-4}$ of 0.6. Due to the network structure, many downstream pathways in this model saturate at an input level of 0.65 as was shown previously[1]. Setting an input's peak to 0.6 indicates that it approaches saturation while still allowing for the potential of further signaling through that pathway. Parameters for each input curve are provided in **Table S3**.

**Equation 1.1:** Single-peak input equation

$$1 + p_1 e^{-\frac{t}{p_2}}(1 - e^{-\frac{t}{p_3}})$$

**Equation 1.2:** Double-peak input equation

$$1 + p_1 e^{-\frac{t}{p_2}}\left(1 - e^{-\frac{t}{p_3}}\right) + p_4 e^{-\frac{t}{p_5}}\left(1 - e^{-\frac{t}{p_6}}\right)$$

#### *Tissue-Level Model*

The single-cell signaling model was coupled to a previously reported model of the tissue-level accumulation of collagen (collagen area fraction) [10]. Whereas the tissue-level model previously used a fixed time-dependent curve for collagen mRNA levels, here we drove the model by the collagen I mRNA dynamics predicted by the fibroblast signaling network model. The change in collagen area fraction was modeled based on **Equation 2**. Specifically, the production of collagen is modeled as a product of the collagen generation rate ($k_g$, 1.8 unit/day), the collagen I mRNA levels ($c_n$ as predicted by the network model), and the number of fibroblasts ($n_f$). Fibroblast number was based on observed changes in fibroblast proliferation post-MI [11]. The degradation of collagen is the product of the degradation rate ($k_d$, 0.03 unit/day), the level of MMP activity (m), and the amount of mature collagen. The MMP activity level was modeled with **Equation 3**, based on the average of the tissue-level dynamics of MMPs 1, 2, and 9[12].

**Equation 2:**

$$\frac{dAreaFraction}{dt} = k_g * c_n * n_f - k_d * m * AreaFraction$$

**Equation 3:**

$$m(t) = kd_1 + kd_2 * \boldsymbol{e}^{(-\boldsymbol{kd_3}*\boldsymbol{t})} - \boldsymbol{e}^{(-\boldsymbol{kd_2}*\boldsymbol{t})}$$

$$kd_1 = 0.2 \quad kd_2 = 0.5592 \quad kd_3 = 0.10368$$

[7] Voloshenyuk TG, Hart AD, Khoutorova E, et al. TNF-?? increases cardiac fibroblast lysyl oxidase expression through TGF-?? and PI3Kinase signaling pathways. *Biochem Biophys Res Commun*; 413: 370–375.

[8] Hall MC, Young DA, Waters JG, et al. The comparative role of activator protein 1 and Smad factors in the regulation of Timp-1 and MMP-1 gene expression by transforming growth factor-β1. *J Biol Chem*; 278: 10304–10313.

[9] Yuan W, Varga J. Transforming growth factor-beta repression of matrix metalloproteinase-1 in dermal fibroblasts involves Smad3. *J Biol Chem*; 276: 38502–10.

[10] Clarke SA, Richardson WJ, Holmes JW. Modifying the mechanics of healing infarcts: Is better the enemy of good? *Journal of Molecular and Cellular Cardiology* 2016. Epub ahead of print 2016. DOI: 10.1016/j.yjmcc.2015.11.028.

[11] Fishbein MC, Maclean D, Maroko PR. The histopathologic evolution of myocardial infarction. *Chest*; 73: 843–849.

[12] Cleutjens JPM, Kandala JC, Guarda E, et al. Regulation of collagen degradation in the rat myocardium after infarction. *J Mol Cell Cardiol* 1995. Epub ahead of print 1995. DOI: 10.1016/S0022-2828(05)82390-9.

### Supplemental Media

Supplemental Video 1: Network visualization of dynamic post-MI model showing time-dependent activation of each network node in response to the dynamic paracrine input curves.

### Supplemental Tables

**Table S1**: Supporting data for additional reactions in model update

| Reaction | Primary source cell type | Primary Source PMID | Secondary source cell type | Secondary Source PMID | Additional PMID |
|---|---|---|---|---|---|
| **Akt => LOX** | rat CF | 21498085 | rat CF | 21893029 | |
| **LOX & CI => xlinkFibers** | human myocardium | 19075089 | mouse lung | 23345161 | |
| **AP1 & !smad3 => proMMP1** | human cardiac fibroblast | 17921324 | human dermal fibroblasts | 11502752 | 12525489 |
| **FA => contraction** | rat dermal fibroblast, rat lung fibroblast | 11553712 | human corneal fibroblast | 17965264 | |
| **aSMA => contraction** | rat dermal fibroblast, rat lung fibroblast | 11553712 | human corneal fibroblast | | |

**Table S2**: Post-Infarct Measurements of Model Inputs

| speciesName | 0 | <1 day peal | 1day | 3day | 7day | 2wk | 4 wk | 6 week | 7 week | Location | citation |
|---|---|---|---|---|---|---|---|---|---|---|---|
| Angiotensin | 20.5518 | 24.60842 | | 28.11324 | | 18.7024 | | | | human serum | 11717612 |
| Angiotensin | 0.27 | | | | | | | | 0.64 | rat whole LV | 8181153 |
| TGFb | 0.047392 | 0.414658 | 0.094737 | | 0.903393 | | | | | rat whole heart | 7856752 |
| TGFb | 0.041558 | 0.316883 | 0.093506 | | 0.264935 | | | | | rat whole heart reperfused | 7856752 |
| TGFb | | 8.487281 | 7.72866 | 4.203505 | 4.566422 | 15.04466 | 25.95025 | | | rat infarct | 11444923 |
| IL6 | 0.014285 | 0.763522 | 0.066663 | | 0.112539 | | | | | rat whole heart | 7856752 |
| | 0.009565 | 0.358523 | 0.023829 | | 0.02078 | | | | | rat whole heart reperfused | 7856752 |
| | 0.625 | 46.5625 | 10.625 | 1.5625 | 1.5625 | 0.62 | 0 | | | rat infarct | 12123772 |
| IL1b | 0.029577 | 0.490141 | 0.088732 | | 0.54507 | | | | | rat whole heart | 7856752 |
| | 0.030435 | 0.56087 | 0.065217 | | 0.082609 | | | | | rat whole heart reperfused | 7856752 |
| | 1.8 | 36.7248 | 19.2607 | 6.4476 | 6.69404 | 7.9671 | 11.817 | | | rat infarct | 12123772 |
| TNFa | 0.075342 | 0.43505 | 0.107747 | | 0.884695 | | | | | rat whole heart | 7856752 |
| | 0.064 | 0.549333 | 0.032 | | 0.165333 | | | | | rat whole heart reperfused | 7856752 |
| | 0.2 | 0.1625 | | | | | 0.23125 | | | rat infarct border | 11399901 |
| NE | 289.3617 | 659.5745 | | 455.319 | | 468.0851 | | | | human serum | 11717612 |
| ET1 | 0.695535 | 1.530866 | | 1.129083 | | 1.120692 | | | | human serum | 11717612 |
| BNP | 7.423295 | 56.66477 | | 70.60227 | | 81.35796 | # | | | human serum | 11717612 |
| PDGF | 1.015 | | 0.4 | 1.092 | 2.708 | 3.138 | 2.323 | 1.308 | | rat border zone | 21767547 |
| | 0.968 | | 0.255 | 2.776 | 3.638 | 4.521 | 2.808 | 2.117 | | rat infarct | 21767547 |

Color indicates whether the value at that time point is a peak. Pink indicates a low peak (<3x change), red indicates a high peak (>3x change), blue indicates a value lower than pre-infarct levels. The citation indicates the PMID of the paper from which the data was collected. All values interpolated from figures in the indicated papers using digitize2.m (https://www.mathworks.com/matlabcentral/fileexchange/928-digitize2-m).

**Table S3**: Parameters for each of the dynamic post-MI paracrine inputs

**Table S3a** One-Peak input curves

| Input | p1 | p2 | p3 |
|---|---|---|---|
| **IL6** | 11216.263 | 10 | 550 |
| **BNP** | 14.953 | 1200 | 160 |
| **AngII** | 2.5587 | 3200 | 750 |
| **PDGF** | 10.902 | 400 | 560 |

**Table S3b** Two-Peak input curves

| Input | p1 | p2 | p3 | p4 | p5 | p6 |
|---|---|---|---|---|---|---|
| **TGFb** | 440.673 | 170 | 1500 | 25 | 2000 | 5900 |
| **IL1** | 40.452 | 18 | 6 | 180 | 750 | 8600 |
| **TNFa** | 2 | 25 | 0.5 | 99.081 | 170 | 540 |
| **NE** | 1.8146 | 10 | 1 | 5 | 220 | 400 |
| **ET1** | 1.6827 | 10 | 1 | 5 | 220 | 400 |

### Supplemental Figures

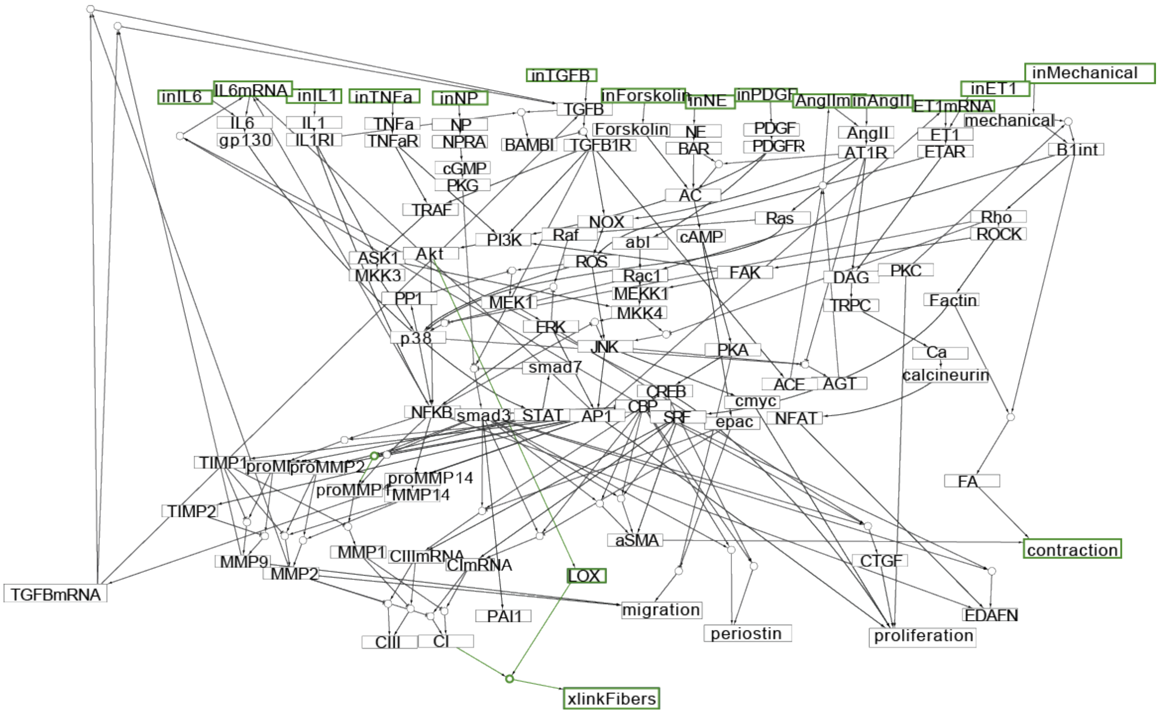

**Figure S1: Network Modifications.** Network diagram of the fibroblast network model, with additions as described in Supplemental Methods. Green color highlights the interactions and nodes that were added to the network model in this study.

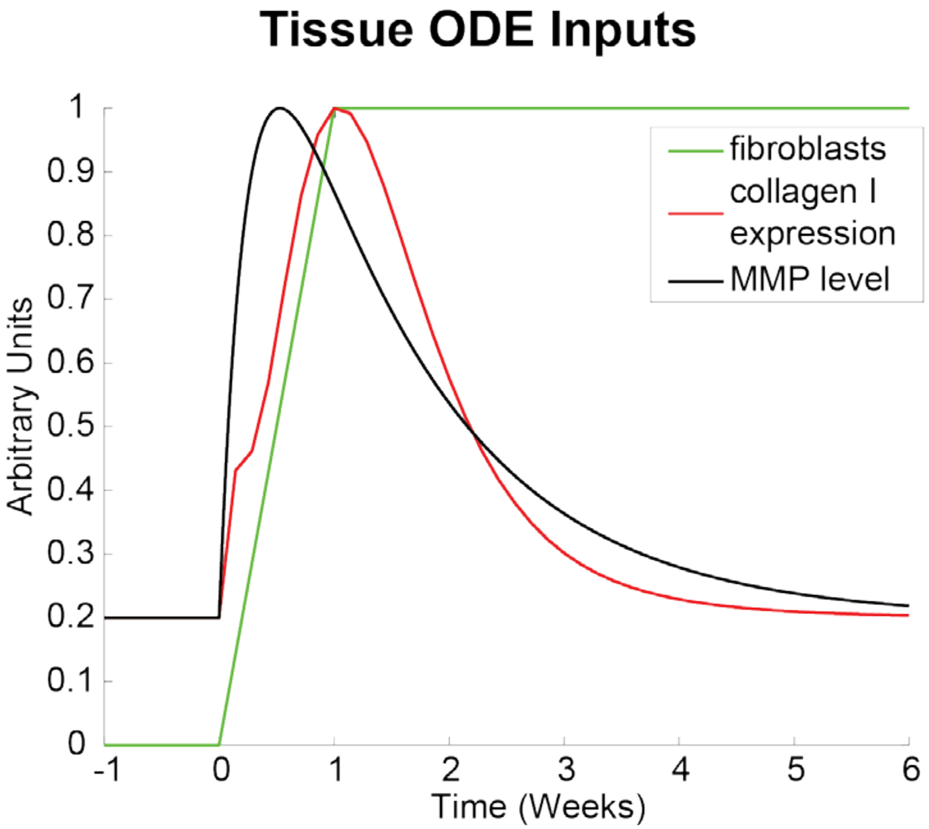

**Figure S2: Inputs to tissue-level model for simulations of post-MI wound healing.** Shown are curves that input to the tissue-level model. Fibroblast number and MMP levels are defined by an idealized input curve based on post-MI data as described in Supplemental Methods. Collagen I expression is the collagen I mRNA level predicted by the network model, normalized to a max value of 1.

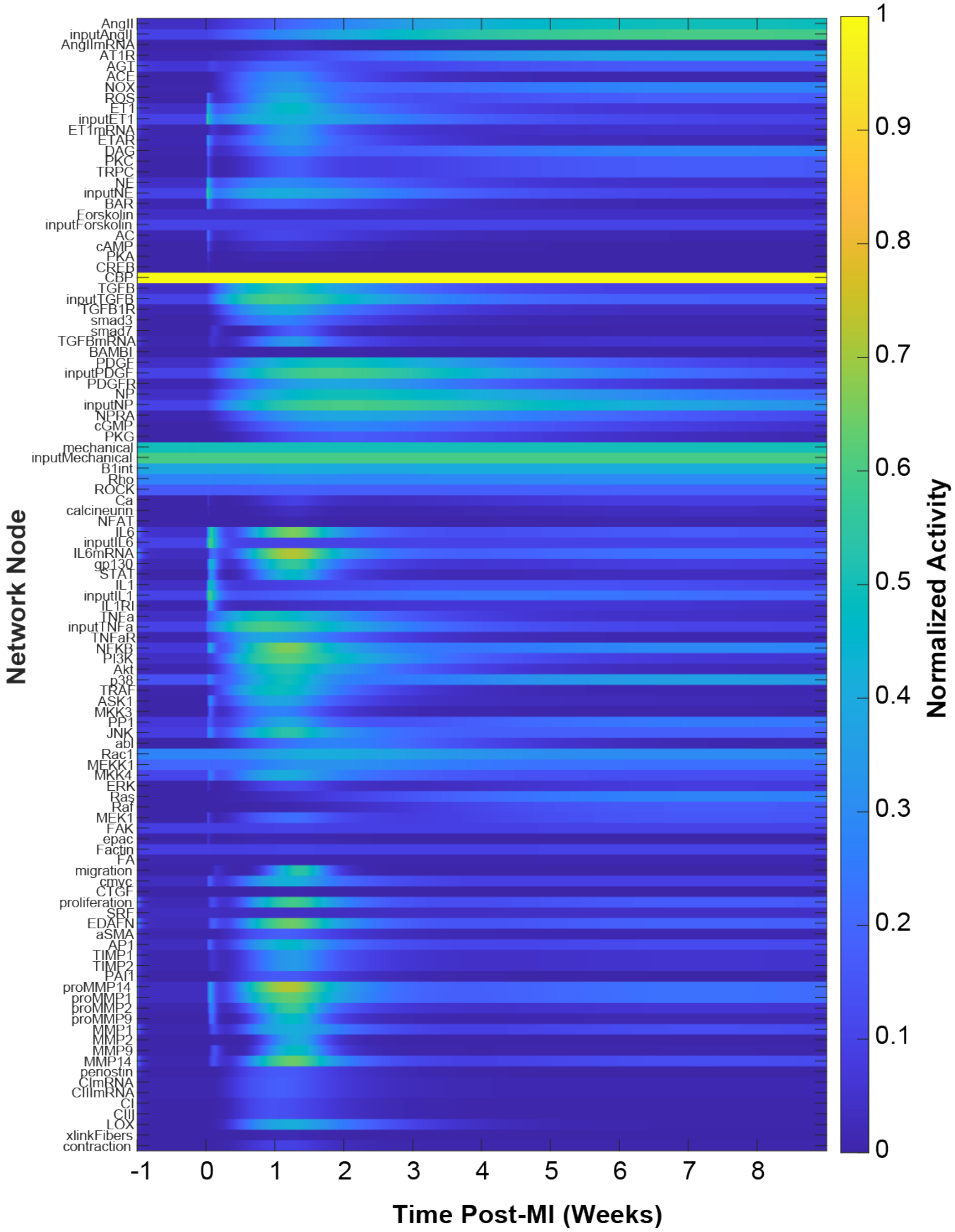

**Figure S3: Predicted dynamics of activation of each network node in simulation following myocardial infarction, with nine dynamic paracrine stimuli.**

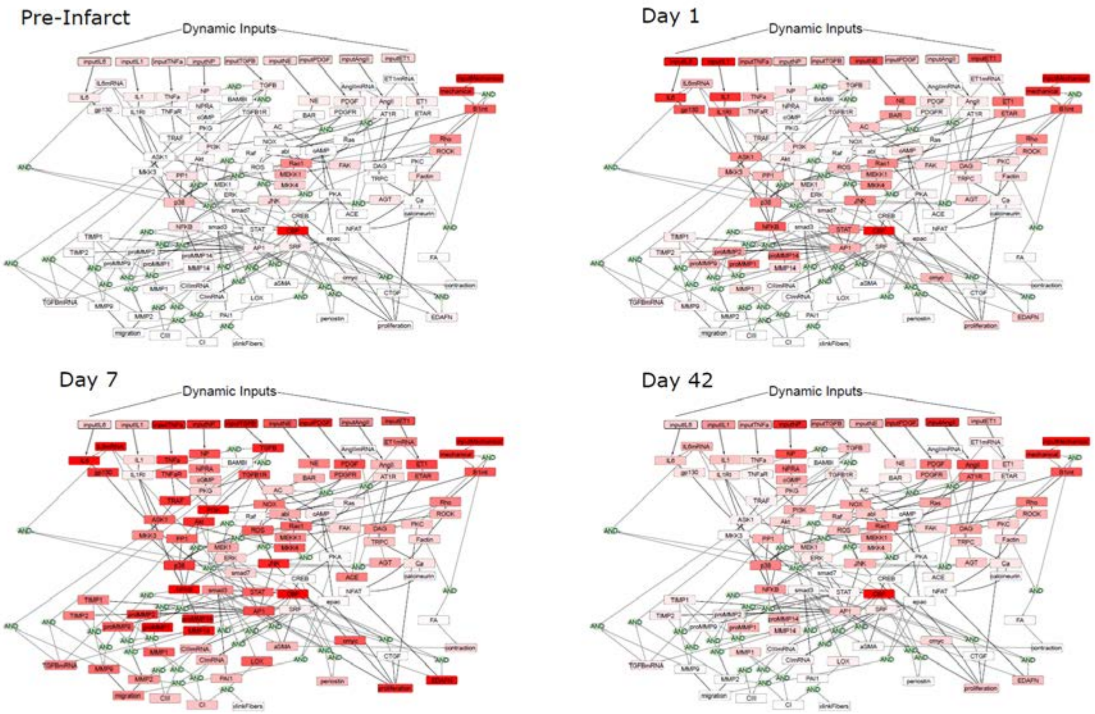

**Figure S4: Network visualization of node activity at key timepoints representative of pre-infarct (Day 0), inflammatory (Day 1), proliferative (Day 7), and maturation (Day 42) phases following myocardial infarction.** This figure visualizes the dynamic post-MI simulation reported in **Figure 2**.

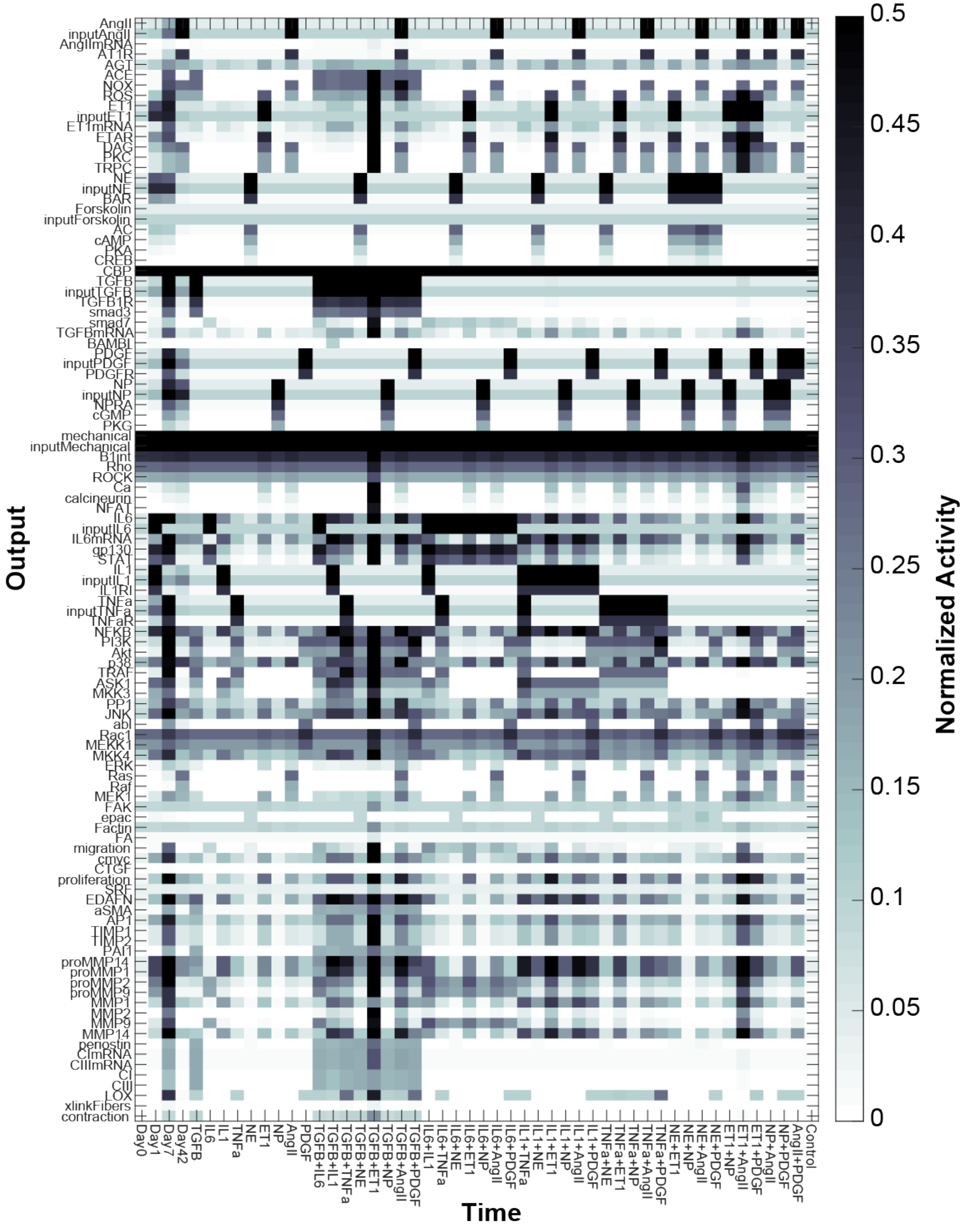

**Figure S5: Network-wide activation profiles in response to each single-input and double-input static condition.** Each row represents a different node, and each column represents a time point in the dynamic post-MI simulation (Days 0-42) or a static simulation with a single static paracrine stimulus, two static paracrine stimuli, or no stimulus (control).

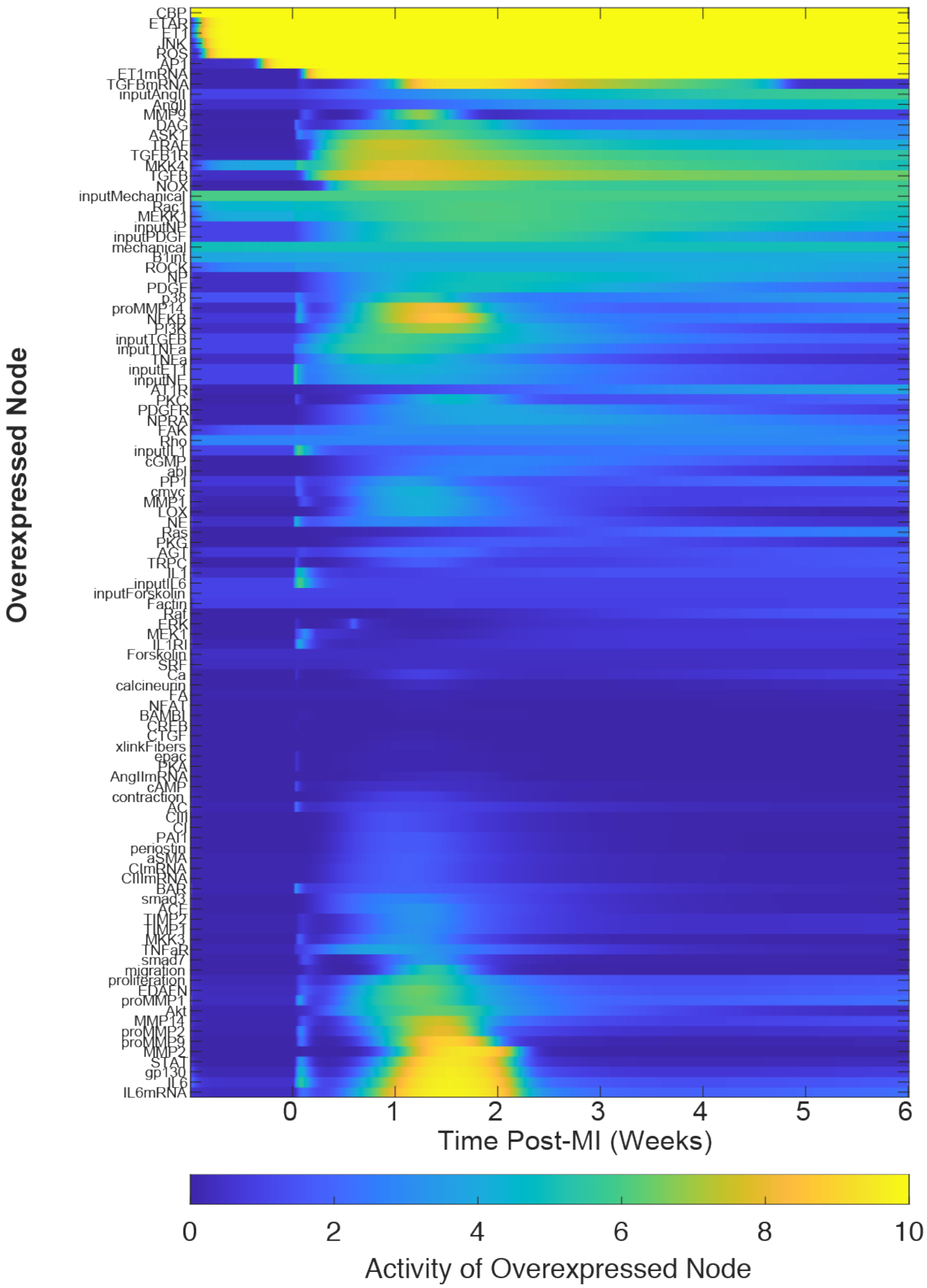

**Figure S6**: Dynamics of overexpressed nodes from the overexpression screen with dynamic post-MI inputs. As in **Figure 5**, each row represents a simulation in which the nine dynamic paracrine stimuli were used as inputs, with the indicated node increased from ymax = 1 to ymax = 10. Here, the heatmap shows the dynamic activity of that overexpressed node.

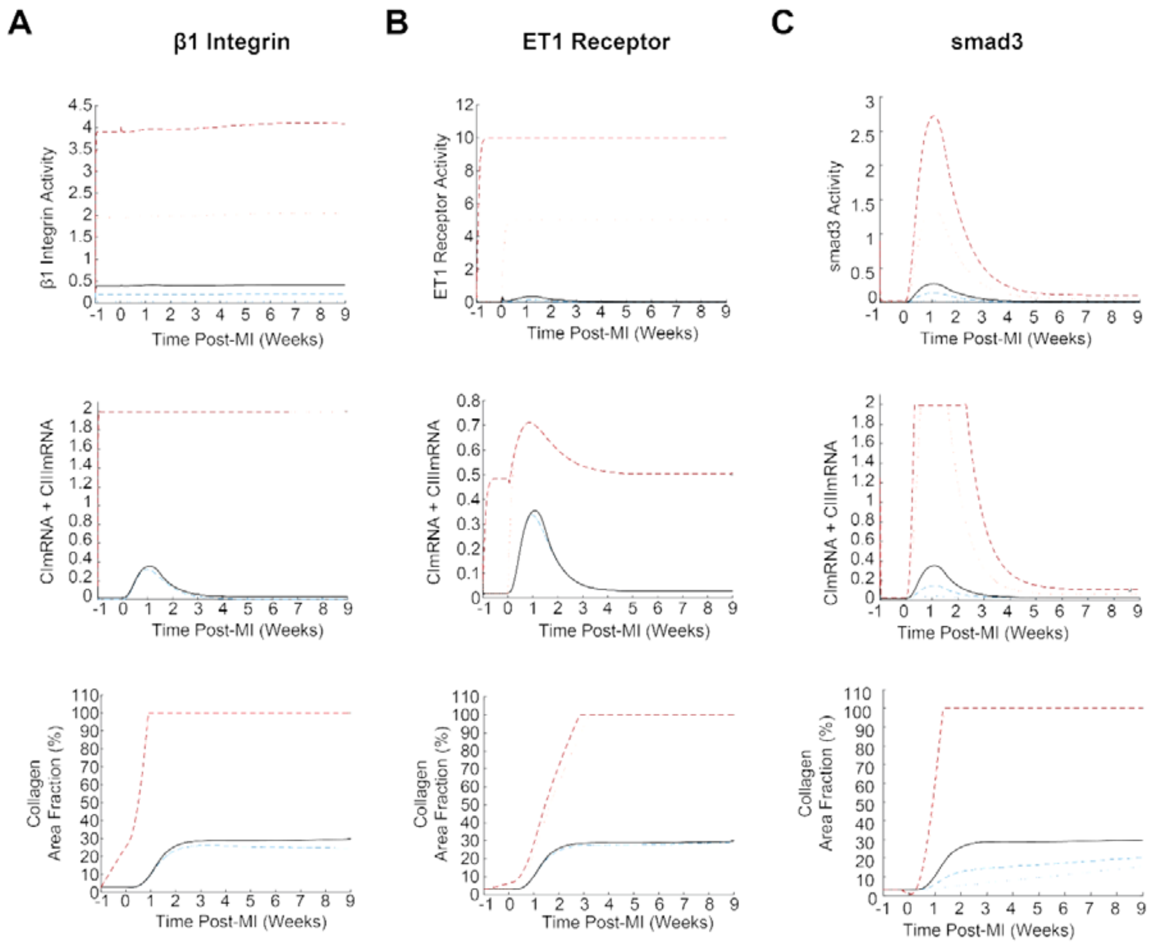

**Figure S7: Mechanisms contributing to regulation of collagen expression post-MI with altered expression of A) β1-integrin, B) ET1AR, and C) smad3.** As in **Figure 6**, node activity, collagen mRNA expression, and collagen area fraction are shown for control levels of expression (black line, ymax = 1), overexpression (red dashed line, ymax = 10), or knockdown (blue dashed line, ymax = 0.1) of the indicated node.
